# tbiExtractor: A framework for extracting traumatic brain injury common data elements from radiology reports

**DOI:** 10.1101/585331

**Authors:** Margaret Mahan, Daniel Rafter, Hannah Casey, Marta Engelking, Tessneem Abdallah, Charles Truwit, Mark Oswood, Uzma Samadani

## Abstract

**Objective:** The manual extraction of valuable data from electronic medical records is cumbersome, error-prone, and inconsistent. By automating extraction in conjunction with standardized terminology, the quality and consistency of data utilized for research and clinical purposes would be substantially improved. Here, we set out to develop and validate a framework to extract pertinent clinical conditions for traumatic brain injury (TBI) from computed tomography (CT) reports.

**Materials and Methods:** We developed tbiExtractor, which extends pyConTextNLP, a regular expression algorithm using negation detection and contextual features, to create a framework for extracting TBI common data elements from radiology reports. The algorithm inputs radiology reports and outputs a structured summary containing 27 clinical findings with their respective annotations. Development and validation of the algorithm was completed using two physician annotators as the gold standard.

**Results:** tbiExtractor displayed high sensitivity (0.92-0.94) and specificity (0.99) when compared to the gold standard. The algorithm also demonstrated a high equivalence (94.6%) with the annotators. A majority of clinical findings (85%) had minimal errors (F1 Score ≥ 0.80). When compared to annotators, tbiExtractor extracted information in significantly less time (0.3 sec vs 1.7 min per report).

**Discussion and Conclusion:** tbiExtractor is a validated algorithm for extraction of TBI common data elements from radiology reports. This automation reduces the time spent to extract structured data and improves the consistency of data extracted. Lastly, tbiExtractor can be used to stratify subjects into groups based on visible damage by partitioning the annotations of the pertinent clinical conditions on a radiology report.

## INTRODUCTION

Radiology reports from electronic medical records (EMR) are formatted as unstructured narrative text meant for human consumption and contain vast amounts of detailed information that is underutilized. For example, one of the most valuable sources of information for assessing traumatic brain injury (TBI) is the initial head computed tomography (CT) scan. Notably, CT findings have been shown to be one of the most powerful prognosticators in assessing six-month outcomes in TBI [1]. However, extracting structured information from radiology reports is time consuming, error-prone, requires trained professionals for accuracy, and is inconsistent across clinical trial sites and research studies [2].

To address these inconsistencies, many fields have adopted common data elements [3], which are predefined units of information to be used in a collaborative fashion; in other words, a set of uniform terminology [4]. By enabling this interoperability of data via common data elements, the design of clinical trials and research studies based on standard stratification of subject groups is possible [2]. It should be noted that two standardized classifications of TBI based on CT findings have been widely used, namely Marshall and Rotterdam scores [5–6]. However, these classifications focus on subjects with severe injuries and forgo granularity in variables describing the underlying pathology. Therefore, the focus on common data elements remains important for detailing TBI injuries.

Even with the adoption of common data elements, extracting structured information from radiology reports is limited by manual annotation, which takes time and is error-prone. Bypassing this human bottleneck through automation has the potential to expedite research findings, aid in large-scale clinical trials, and ultimately improve clinical care for TBI patients [7]. To facilitate this automation, natural language processing methods can be utilized to parse free-text clinical narratives from EMRs by analyzing linguistic concepts and categorizing them appropriately [8–9].

The field of natural language processing is extensive with a diverse set of subproblems [10] that have been implemented in a variety of medical contexts [11–21]. Four subproblems of interest are problem-specific segmentation, named entity recognition, negation and uncertainty identification, and information extraction. Problem-specific segmentation aims to separate text into groups; for example, segmenting sections of a radiology report into “History” and “Findings” sections. Named entity recognition aims to identify and categorize specific words or phrases; for example, categorizing a set of radiology reports based on type of scan (e.g., head CT vs lumbar spine CT). Negation and uncertainty identification aim to identify specific words or phrases as present or absent; for example, “no evidence of intracranial pathology” would indicate absence of pathology. Information extraction aims to identify and translate problem-specific information into structured data; for example, identifying mass lesions on a radiology report requiring surgical evacuation.

## OBJECTIVE

The purpose of our study was to develop and validate an algorithm, termed tbiExtractor, which incorporates natural language processing methods to extract twenty-seven common data elements from radiology reports in an automated fashion. The output provides a structured summary of pertinent clinical conditions for a TBI subject. Successful implementation of this algorithm has the potential to reduce the time spent to extract structured data, improve the quality of data extracted, and provide a mechanism for systematic placement of subjects into research groups.

## MATERIALS AND METHODS

Development and analysis were performed using Python 3.6.6 [22] with the following libraries: Pandas (0.23.4) [23], NumPy (1.15.0) [24], SciPy (1.1.0) [25], spaCy (2.0.12) [26], scikit-learn (0.19.2) [27], pyConTextNLP (0.6.2.0) [28], NetworkX (1.11) [29], Matplotlib (2.2.3) [30], and Seaborn (0.9.0) [31]. A methods flowchart is shown in Fig 1.

**Fig 1.**
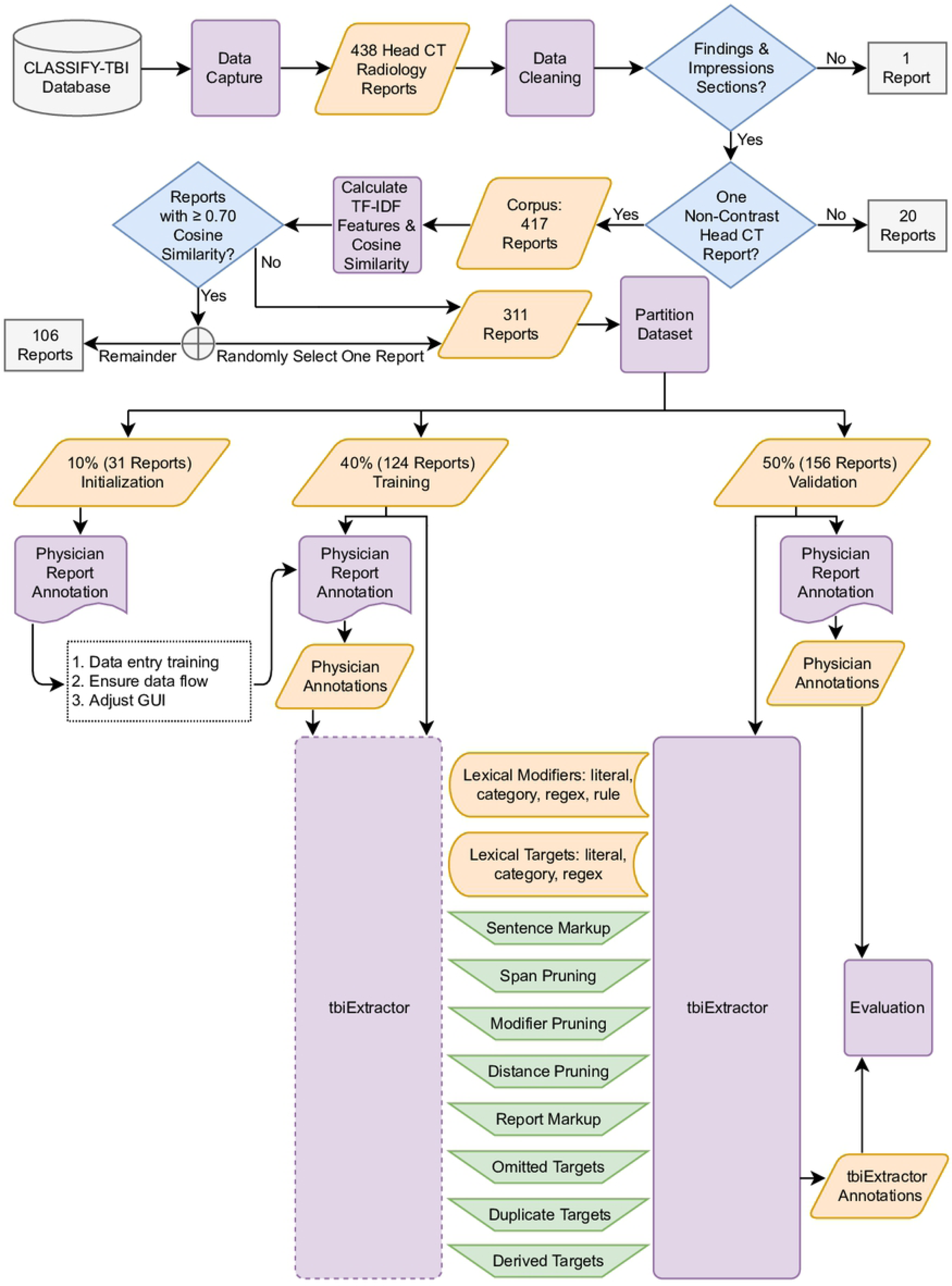
Graphical outline of the methods. Purple rectangle shapes correspond to methods subsections, meaning they represent steps in the processing workflow, orange parallelogram shapes represent data, blue diamond shapes represent binary decisions on data, gray rectangle shapes represent excluded data, and green isosceles trapezoid shapes correspond to subcomponents of the algorithm.

### Data Capture and Cleaning

Hospital admission radiology reports from non-contrast head CT scans were extracted from EMRs for subjects participating in the CLASSIFY-TBI study (details in S1 Protocol). Each radiology report was converted to a spaCy [26] container for assessing linguistic annotations and partitioned into sentences. Sentences before “Findings” and after “Impressions” sections were removed. Then, the sentences were concatenated with newline characters replaced with a space, symbols removed, and whitespace stripped. Radiology reports that did not contain “Findings” or “Impressions” sections were removed along with radiology reports containing multiple scan types.

### Calculate TF-IDF and Cosine Similarities

Using scikit-learn [27] TfidfVectorizer, the corpus was converted into a matrix of TF-IDF (term-frequency times inverse document-frequency) features using *n*-grams with *n*-range from one to ten. Cosine similarities were calculated between each pair of radiology reports by multiplying the TF-IDF matrix by its transpose. Using the cosine similarity for each pair of radiology reports, one radiology report was randomly selected and all radiology reports with at least 0.70 cosine similarity to that radiology report were collected in a set. From this set, one radiology report was randomly selected to keep for further analysis and the remainder were removed. This was applied recursively for each set until each radiology report was retained for further analysis or marked for removal. The purpose of this removal was to reduce the data requiring human annotation. Details in S2 Appendix.

### Dataset Partitioning

A random deck of three numbers the same size as the number of radiology reports retained for analysis was created. The three numbers represented the proportion of radiology reports to be assigned to each of the datasets: 10% initialization, 40% training, and 50% validation. From the set of radiology reports retained for analysis, one radiology report was randomly selected along with up to three most similar radiology reports, based on cosine similarity. From this subset, each radiology report was assigned the next number in the shuffled deck. This was applied recursively until each radiology report was assigned to one dataset.

The initialization dataset was solely used for training annotators and was not used by the algorithm, the training dataset was used to enhance the development of the algorithm by incorporating input from annotators, and the validation dataset was used to compare the annotators to the developed algorithm to determine the algorithm’s viability.

### Radiology Report Annotation

A custom-built Graphical User Interface (GUI) was developed using Python’s TkInter library [22]. The GUI presented two physician annotators with one de-identified radiology report and drop-down menus, one for each lexical target with the respective annotation options. Annotators viewed one radiology report at a time and were not allowed to edit their annotations after submission. The annotation options were: PRESENT, SUSPECTED, INDETERMINATE, NOT SPECIFIED, ABSENT, NORMAL, ABNORMAL (definitions in S3 Appendix). Each dataset was presented to annotators separately. The initialization dataset was used to train annotators on the data entry process and scope of the project, ensure the data processing flow was valid, and make adjustments to the GUI. The training and validation datasets were presented to the annotators and their annotations were retained for development and validation of the algorithm, respectively.

### tbiExtractor Development

We developed tbiExtractor, which extends pyConTextNLP [28] to create a framework for extracting TBI common data elements from radiology reports [3]. tbiExtractor inputs a non-contrast head CT radiology report and outputs a structured summary containing 27 common data elements with their respective annotations. For example, subdural hemorrhage (common data element) is PRESENT (annotation). Code and data files to implement tbiExtractor, along with a Jupyter notebook tutorial, are available at https://github.com/margaretmahan/tbiExtractor.

#### pyConTextNLP Background

Based on a regular expression algorithm called NegEx [32], which uses negation detection (e.g., no evidence of intracranial pathology), the ConText [33–34] algorithm captures the contextual features surrounding the clinical condition by relying on trigger terms and termination clues. A more extensible version of the ConText algorithm was implemented in Python, pyConTextNLP [28], and offers added flexibility for user-defined contextual features and indexed events (e.g., specific clinical conditions) [35].

As a lexicon-based method, pyConTextNLP inputs tab-separated files for lexical targets (indexed events) and lexical modifiers (contextual features). It then converts these into itemData, which contains a literal, category, regular expression, and rule (the latter two are optional). The literal, belonging to a category (e.g., ABSENT), is the lexical phrase (e.g., is negative) in the text. The regular expression allows for variant text phrases (e.g., was negative) giving rise to the same literal and is generated from the literal if not provided. Further, the rule provides context to the span of the literal (e.g., backward).

For text data, pyConTextNLP marks the text with lexical modifiers and lexical targets according to their representative itemData. The pyConTextNLP algorithm outputs a directional graph via NetworkX[29] which represents these markups. Nodes in the graph represent the concepts (i.e., lexical modifiers and lexical targets) in the text and edges in the graph represent the relationship between the concepts.

The following three subsections will describe the details used for extending pyConTextNLP.

#### Lexical Modifiers and Lexical Targets

Lexical modifiers were adapted from a pyConTextNLP application to CT pulmonary angiography reports[35]. Modifications in deriving the final lexical modifiers are as follows:

1. The literal is a lexical phrase (e.g., was not excluded). Literals were added and removed during the training stage.
2. The category is what the literal refers to (e.g., INDETERMINATE). Each literal was assigned a category before the initialization stage and updated during the training stage. The categories used for this study are PRESENT, SUSPECTED, INDETERMINATE, NOT SPECIFIED, ABSENT, NORMAL, and ABNORMAL. Henceforth, the term “annotation” will be used when referencing the category to maintain consistency between annotators and algorithm vocabulary.
3. The regular expression is used to find variant text phrases (or patterns) for the same literal (e.g., the regular expression: `(was|were)\snot\sexcluded`, would find sentences with “was not excluded” and “were not excluded”). Regular expressions were added and updated during the training stage.
4. The rule dictates the span of the literal (e.g., backward). Each literal was assigned a rule before the initialization stage and updated during the training stage. The rules used for this study are forward, backward, and bidirectional.

Lexical targets were adapted from the common data elements in radiologic imaging of TBI [3]. In deriving the lexical targets, the literal represents a clinical condition relevant to TBI on a non-contrast head CT scan (e.g., microhemorrhage) and the category, in this study, is the same (e.g., MICROHEMORRHAGE). The regular expression for each literal (e.g., `microhemorrhage(s)?`) was added and updated during the training stage.

Two examples (Fig 2 and Fig 3) are provided for detailed explanation of the application of lexical modifiers and lexical targets during the algorithm process.

**Fig 2.**
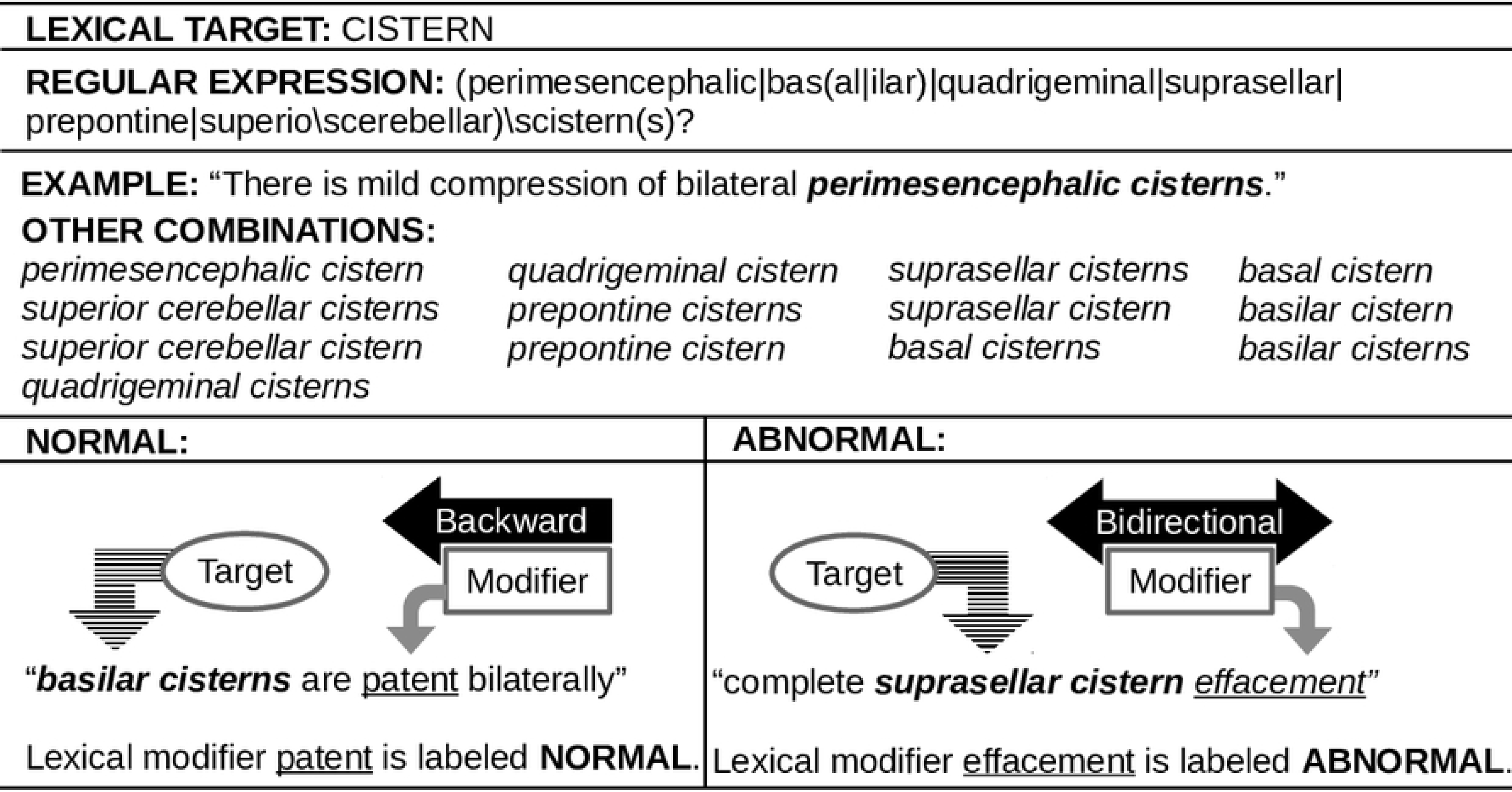
First example of the application of lexical targets and lexical modifiers.

**Fig 3.**
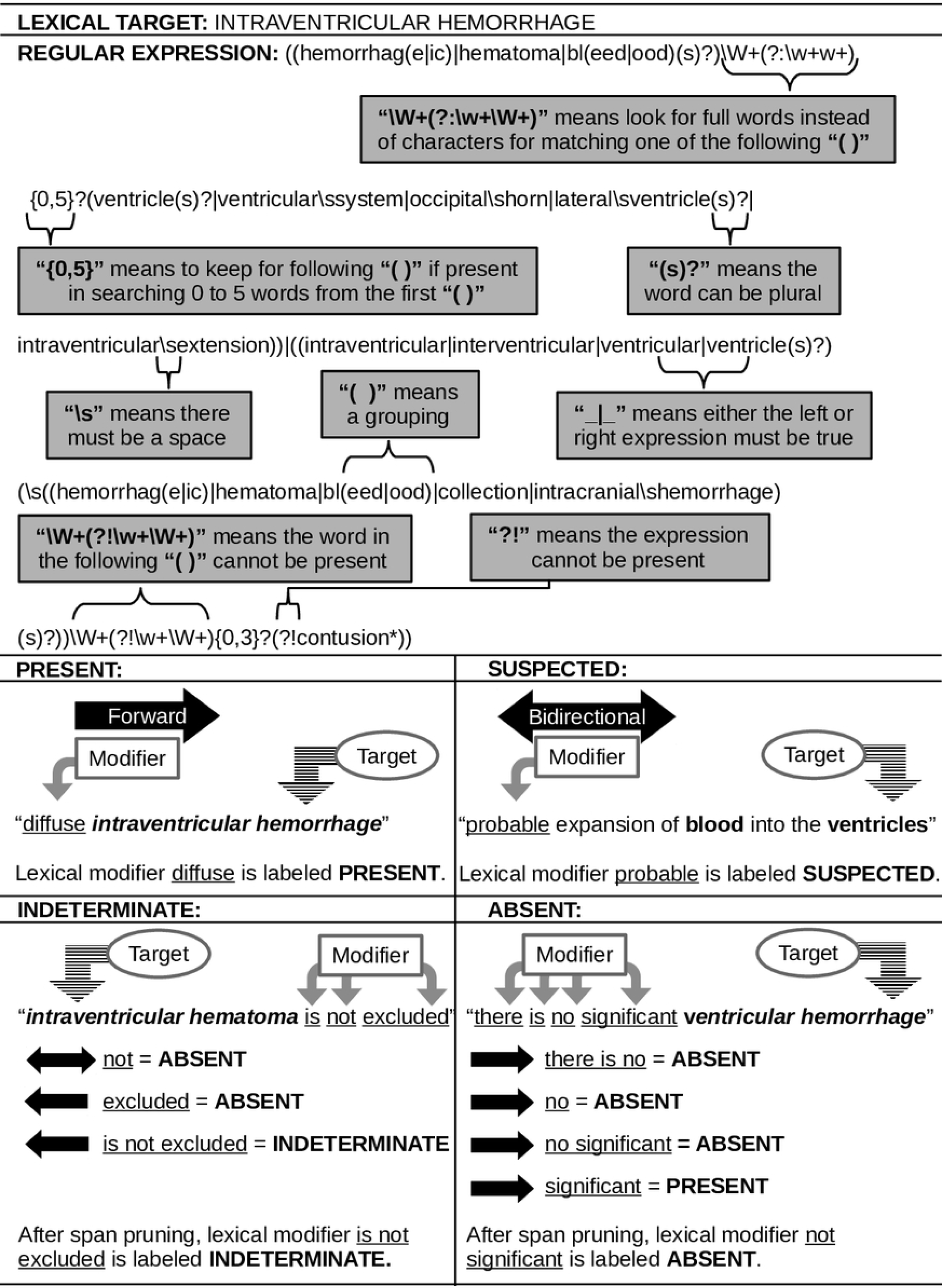
Second example of the application of lexical targets and lexical modifiers.

#### Sentence Markup Followed by Span, Modifier, and Distance Pruning

To implement tbiExtractor, each cleaned radiology report was converted to a spaCy [26] container and subsequently partitioned into sentences. Using pyConTextNLP [28], each sentence was marked with lexical modifiers and lexical targets according to their representative itemData. Following the markup, concepts that are a subset of another concept, within the same concept type, are pruned (span pruning). For example, if the text contained the phrase “findings do not appear significantly changed”, the lexical modifier <u>not</u> would be pruned and the lexical modifier <u>do not appear significantly changed</u> would be retained. Then, for the marked lexical targets, the lexical modifiers are applied. Lexical modifiers that are not linked to a lexical target are dropped (modifier pruning). For multiple lexical modifiers for the same lexical target in the same sentence, the nearest lexical modifier by character length is chosen (distance pruning). For example, if the text contained the phrase “multifocal subarachnoid hemorrhage as described above most notably in the right sylvian fissure”, the lexical modifier <u>multifocal</u> would be selected via distance pruning over the lexical modifier <u>in the</u> since it is closer in character length to the lexical target, *subarachnoid hemorrhage*. Span and modifier pruning are part of the pyConTextNLP implementation. Distance pruning was added as part of tbiExtractor.

At this stage of processing, each sentence in the radiology report will be marked with lexical targets and linked lexical modifiers. There will be one lexical modifier assigned to one lexical target.

#### Report Markup with Revisions for Omitted, Duplicate, and Derived Targets

A radiology report may have duplicate lexical targets if identified in multiple sentences within the radiology report or a radiology report may be lacking lexical targets. To mitigate this, tbiExtractor employs decision rules. First, for each radiology report, omitted lexical targets are added with the default annotation of NORMAL for *gray-white matter differentiation* and *cistern* lexical targets and annotation of ABSENT for the remaining 25 lexical targets (omitted targets). Second, if duplicate lexical targets are identified, the majority vote is selected (duplicate targets). For example, if a lexical target appears in the radiology report three times and the lexical modifiers for two occurrences have an annotation of ABSENT and the other has an annotation of PRESENT, tbiExtractor will choose ABSENT. Similarly, if there are two lexical modifiers with an annotation of PRESENT, two with ABSENT, and one with SUSPECTED, tbiExtractor removes SUSPECTED based on the majority vote. However, the annotations PRESENT and ABSENT require further decision rules because no majority exists.

In the case where no majority exists, the first lexical modifier in the ordered annotation list is selected. If the lexical target is *extraaxial fluid collection, hemorrhage not otherwise specified (NOS)*, or *intracranial pathology*, the ordered annotation list is: ABSENT, INDETERMINATE, SUSPECTED, PRESENT, NORMAL, ABNORMAL. For all other lexical targets, the ordered annotation list is: PRESENT, SUSPECTED, INDETERMINATE, ABSENT, ABNORMAL, NORMAL. Following this, annotations that are not in the set of annotations for that lexical target are replaced with their predetermined counterpart (e.g., if the lexical target *cisterns* has an annotation of ABSENT, the annotation is replaced with NORMAL). At this stage of processing, each lexical target has one annotation for the entire radiology report.

The annotations for three lexical targets can be altered based on the annotations of other lexical targets in the same radiology report. Thus, a second set of derived decision rules are applied by tbiExtractor (derived targets). First, if *epidural hemorrhage, subdural hemorrhage*, or *subarachnoid hemorrhage*, are PRESENT or SUSPECTED, *hemorrhage (NOS)* is annotated ABSENT. Second, if *epidural hemorrhage, subdural hemorrhage*, or *subarachnoid hemorrhage*, are PRESENT, *extraaxial fluid collection* is annotated PRESENT. If these lexical targets were annotated SUSPECTED, and *extraaxial fluid collection* was annotated ABSENT by default, then *extraaxial fluid collection* is annotated SUSPECTED. If *gray-white differentiation, cistern, hydrocephalus, pneumocephalus, extraaxial fluid collection, midline shift, mass effect, diffuse axonal injury, anoxic, herniation, aneurysm, contusion, brain swelling, ischemia, hemorrhage (NOS), intraventricular hemorrhage*, or *intraventricular hemorrhage* are annotated PRESENT, SUSPECTED, or ABNORMAL, then *intracranial pathology* is annotated PRESENT.

Omitted, duplicate, and derived targets were implemented as part of the tbiExtractor. At the end of the above processing steps, each radiology report will have a list of 27 lexical targets each with one annotation, which constitutes the structured summary output.

### Evaluation

Radiology reports were assessed using standard statistical measures. Annotator reliability was measured using Cohen’s kappa (κ) [36–37]. tbiExtractor was evaluated using standard classification performance metrics (equations in S4 Appendix). True positives were defined as the number of times a lexical target was annotated as PRESENT or ABNORMAL by tbiExtractor and annotators, in the first case. In the second case, SUSPECTED was also assigned to the positive group. True negatives were defined as the number of times a lexical target was annotated as ABSENT or NORMAL by tbiExtractor and annotators. In addition, false positives and false negatives were examined to explore why tbiExtractor errors occurred.

## RESULTS

### Radiology Report Characteristics

There were 438 radiology reports extracted: 1 was removed because it did not have both “Findings” and “Impressions” sections, 20 were removed because they contained more than one scan type, and 106 were removed for high cosine similarity. The remaining 311 reports were split into initialization, training, and validation datasets (Table 1).

**Table 1:** Radiology report characteristics for each of the four subsets of data. Values displayed as minimum, maximum, mean ± standard deviation.

| Dataset | Initialization | Training | Validation | Remainder |
| --- | --- | --- | --- | --- |
| Reports | 31 | 124 | 156 | 106 |
| Sentences | 8, 22<br>$14.5 \pm 4.1$ | 4, 72<br>$14.9 \pm 7.6$ | 6, 32<br>$14.3 \pm 5.1$ | 6, 28<br>$8.9 \pm 3.4$ |
| Words | 58, 268<br>$123.5 \pm 54.1$ | 22, 656<br>$122.5 \pm 75.9$ | 35, 284<br>$121.1 \pm 53.6$ | 43, 315<br>$69.7 \pm 35.9$ |
| Cosine Similarity | 0.34, 0.70<br>$0.47 \pm 0.05$ | 0.14, 0.70<br>$0.44 \pm 0.08$ | 0.26, 0.69<br>$0.46 \pm 0.06$ | 0.35, 1.00<br>$0.69 \pm 0.13$ |

### Analysis of Annotators

In the training dataset, annotators took an average of 2.84 minutes per radiology report. Between 15% and 16% of annotations across radiology reports were selected from default (Table 2). There was high equivalence in annotations between the annotators (N = 3175). Further, there were an additional 424 similar annotations (i.e., one annotation PRESENT and the other SUSPECTED). In contrast, there were only 88 divergent annotations (i.e., one annotation ABSENT or NORMAL and the other PRESENT or ABNORMAL). Overall, the two annotators were in high agreement (κ = 0.861). After training, NOT SPECIFIED was removed as an annotation option secondary to the overlap with ABSENT and INDETERMINATE.

**Table 2:** Number and type of annotation selected for each annotator along with comparisons between them for training and validation sets. There were 3348 possible annotations for training and 4212 for validation.

|  |  | Training | Validation |
| --- | --- | --- | --- |
| Annotator 1 | Absent | 2623 | 3244 |
|  | Present | 405 | 569 |
|  | Suspected | 50 | 74 |
|  | Indeterminate | 22 | 13 |
|  | Abnormal | 21 | 23 |
|  | Normal | 227 | 289 |
|  | Not Specified | 0 | 0 |
| Annotator 2 | Absent | 2593 | 3249 |
|  | Present | 440 | 598 |
|  | Suspected | 56 | 48 |
|  | Indeterminate | 9 | 5 |
|  | Abnormal | 17 | 25 |
|  | Normal | 231 | 287 |
|  | Not Specified | 2 | 0 |
| Comparing Annotators | Equivalent | 94.8% | 96.7% |
|  | Similar | 12.7% | 14.2% |
|  | Divergent | 2.6% | 2.1% |
|  | Cohen's kappa | 0.861 | 0.913 |

In the validation dataset, annotators took an average of 1.67 minutes per radiology report. Similar to the training dataset, 16% of annotations across radiology reports were selected from default (Table 2). For the validation dataset, there was high equivalence in annotations between the annotators (N = 4072), with an additional 598 similar annotations, and only 87 divergent annotations. Overall, the two annotators were in high agreement (κ = 0.913).

### tbiExtractor Performance

tbiExtractor took an average of 0.294 seconds per radiology report. A diagram showing the set of annotations across tbiExtractor and annotators for the validation dataset is shown in Fig 4. When comparing tbiExtractor to annotators, there was high equivalence (N = 3984) and low disagreement (N = 13). For the purposes of evaluating the performance of tbiExtractor, cases where annotators were equivalent was considered the gold standard (Fig 4 dashed line). The evaluation revealed high performance across all metrics (Table 3).

**Table 3:** tbiExtractor performance metrics for two cases of positive selection. In both cases, negative if {absent, normal}.

| Metrics | Case 1:<br>where positive if {present,<br>abnormal} | Case 2:<br>where positive if {present,<br>suspected, abnormal} |
| --- | --- | --- |
| Sensitivity | 0.938 | 0.924 |
| Specificity | 0.993 | 0.993 |
| Positive Predictive Value | 0.957 | 0.960 |
| Negative Predictive Value | 0.990 | 0.987 |
| Accuracy | 0.986 | 0.983 |
| F1 Score | 0.948 | 0.941 |

**Fig 4.**
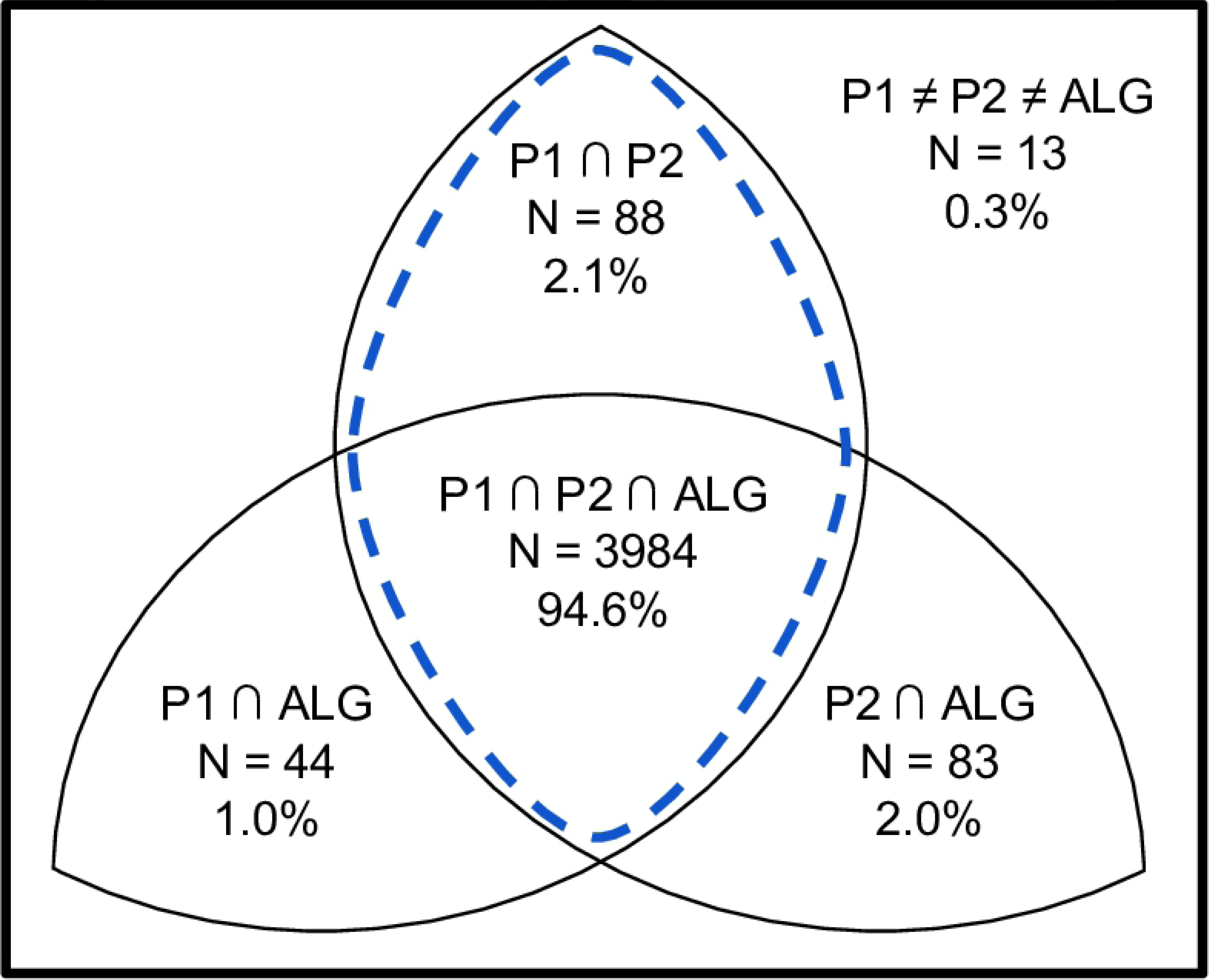
Diagram depicting the overlap in annotations for two annotators (P1, P2) and tbiExtractor (ALG) in validation dataset. Dashed line indicates gold standard (i.e., where two annotators are in agreement).

From the validation dataset (N = 156), the number of lexical targets with equivalence ranged from 20 to 27 (25.5 ± 1.7, mean ± standard deviation), indicating most radiology reports had few errors. Approximately 77% (N = 120) of radiology reports exhibited partial equivalence with at least 25 lexical targets accurately annotated and 93% (N = 145) with at least 23 lexical targets. To show the final lexical targets and their annotations, tbiExtractor was run on the corpus and results for each lexical target are shown in Fig 5.

**Fig 5.**
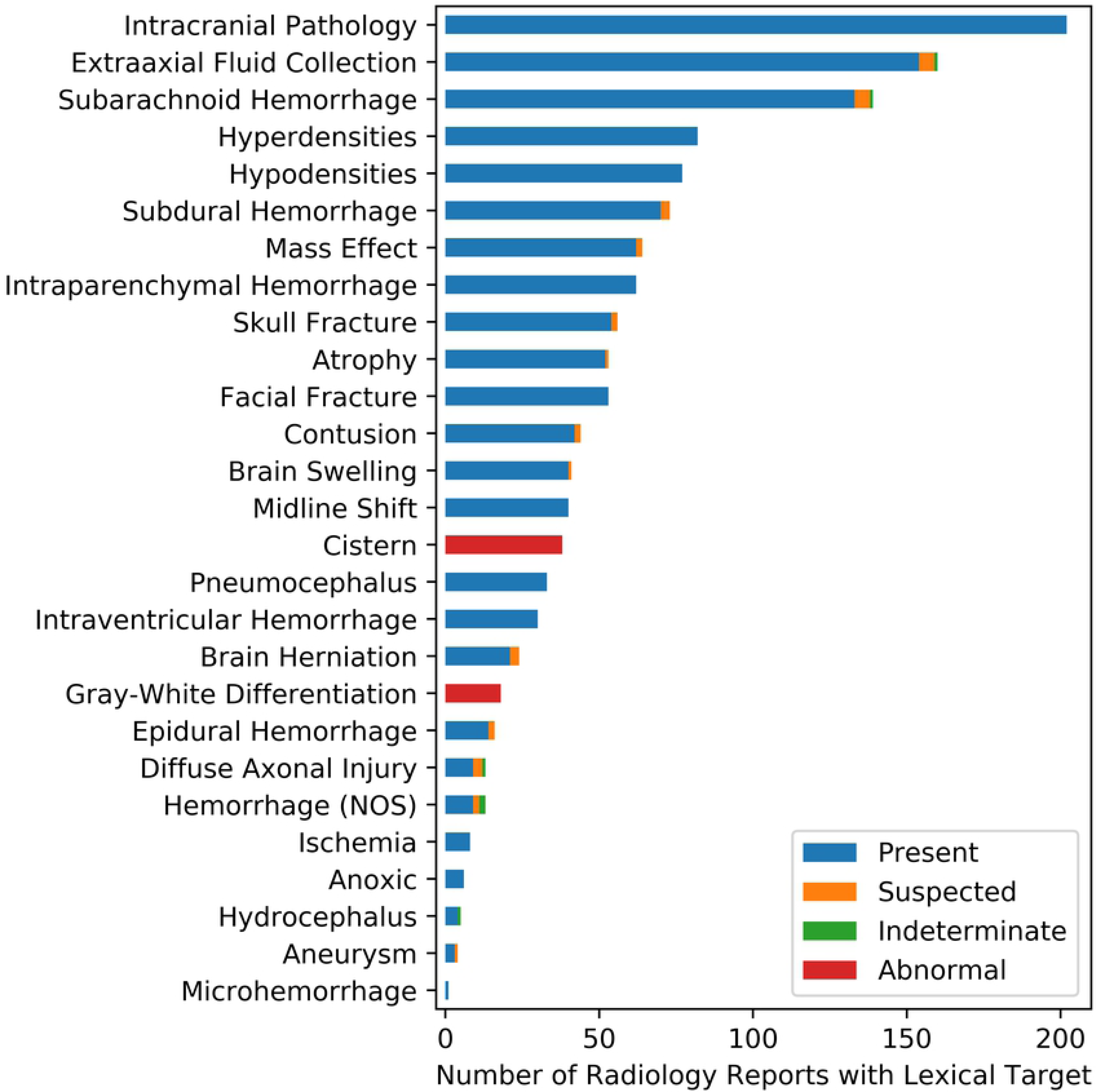
tbiExtractor output annotations for the 27 selected lexical targets over the entire corpus (N = 417 radiology reports).

### Error Analysis

tbiExtractor was evaluated for accuracy on the 27 lexical targets (Table 4). Two lexical targets, *intraparenchymal hemorrhage* and *facial fracture,* produced the most false negatives, meaning tbiExtractor missed these lexical targets outright. This is likely due to the complexity of these lexical targets and the restriction in the regular expressions to term distance (i.e., the distance between fracture and a term indicating facial is more than the allocated {0, 5} from the regular expression). The remaining lexical targets produced minimal false negatives.

**Table 4:** For each of the 27 lexical targets, the occurrences in the validation dataset of each lexical target are displayed with the number of false negatives (FN), false positives (FP), and F1 score performance metric.

| Lexical Target | Occurrences | FN | FP | F1 score |
| --- | --- | --- | --- | --- |
| Intraparenchymal Hemorrhage | 29 | 7 | 1 | 0.85 |
| Facial Fracture | 30 | 6 | 1 | 0.87 |
| Extraaxial Fluid Collection | 75 | 3 | 0 | 0.98 |
| Hypodensities | 36 | 3 | 0 | 0.96 |
| Skull Fracture | 27 | 3 | 0 | 0.94 |
| Intraventricular Hemorrhage | 17 | 3 | 0 | 0.90 |
| Herniation | 11 | 2 | 3 | 0.78 |
| Mass Effect | 26 | 2 | 2 | 0.92 |
| Subarachnoid Hemorrhage | 69 | 2 | 0 | 0.99 |
| Subdural Hemorrhage | 40 | 2 | 0 | 0.97 |
| Hyperdensities | 33 | 2 | 0 | 0.97 |
| Atrophy | 22 | 2 | 0 | 0.95 |
| Contusion | 17 | 2 | 0 | 0.94 |
| Hemorrhage | 2 | 1 | 4 | 0.29 |
| Swelling | 20 | 1 | 2 | 0.93 |
| Pneumocephalus | 17 | 1 | 1 | 0.94 |
| Ischemia | 2 | 1 | 1 | 0.50 |
| Epidural Hemorrhage | 8 | 1 | 0 | 0.93 |
| Anoxic | 4 | 1 | 0 | 0.86 |
| Aneurysm | 3 | 1 | 0 | 0.80 |
| Hydrocephalus | 2 | 1 | 0 | 0.67 |
| Intracranial Pathology | 84 | 0 | 6 | 0.97 |
| Gray-White Differentiation | 7 | 0 | 2 | 0.88 |
| Cistern | 11 | 0 | 1 | 0.96 |
| Midline Shift | 20 | 0 | 0 | 1.00 |
| Diffuse Axonal Injury | 3 | 0 | 0 | 1.00 |
| Microhemorrhage | 1 | 0 | 0 | 1.00 |

Six false positives were produced for *intracranial pathology* and four for *hemorrhage (NOS)*, meaning tbiExtractor identified these lexical targets as PRESENT, while the annotators marked these as ABSENT. This is due to the derivation of these lexical targets in relation to other lexical targets (i.e., if *extraaxial fluid collection* is PRESENT, then by decision rules, so is *intracranial pathology*). The remaining lexical targets produced less minimal false positives. Overall, the errors are minimal as measured by the high F1 scores for the majority of lexical targets.

Further examination of divergent cases (i.e., annotators annotated ABSENT and tbiExtractor annotated PRESENT, or vice versa) revealed the most common diverged lexical targets to be *intracranial pathology, facial fracture, intraparenchymal hemorrhage, hemorrhage (NOS)*, and *herniation*. The remaining lexical targets exhibited less than four diverged responses. The most common lexical modifiers in the divergent cases were the <u>default</u> selection and the derived-from-decision-rules *intracranial pathology*, indicating that most errors were from tbiExtractor missing the lexical targets outright. In most divergent cases where this was not the reason, there were more complex structures to the sentences. In a few other instances, there were sentences that only displayed the lexical target with no available lexical modifier (e.g., *hemorrhagic* extension into the *lateral ventricles*). Two divergent examples are shown in Fig 6.

**Fig 6.**
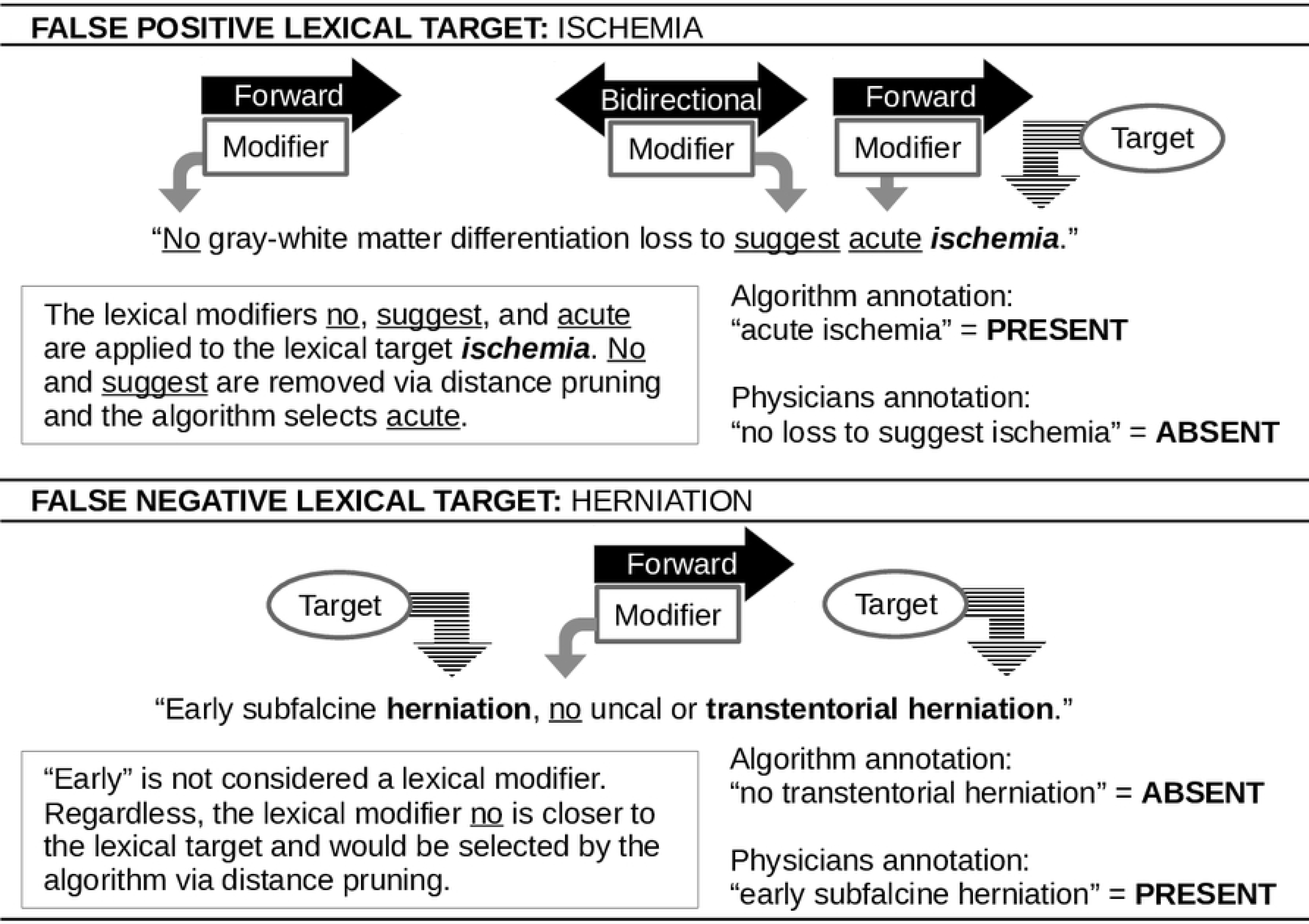
Two examples of divergent annotations between annotators and tbiExtractor.

## DISCUSSION

Assessing the corpus revealed noteworthy characteristics of radiology reports from TBI subjects. First, cosine similarities across the four subsets of data were not different and indicated a normal, albeit slender, distribution of radiology report similarity. Second, the average number of sentences in each radiology reports approached the minimum, indicating a skewed right distribution where the majority of radiology reports will have low numbers of sentences. The same holds true for the number of words. Taken together, this could be reflective of the findings generally found in CT reports on TBI subjects, where the prevalence of CT findings is less than 10% in mild TBI cases [38–42], which constitute approximately 80% of TBI subjects [43–44].

Annotators displayed high equivalent and low divergent annotations. This provided a solid foundation for developing and validating tbiExtractor. In cases where annotators were not equivalent, data entry issues tended to be the culprit. Mostly, this was a result of overlooking the lexical target and not selecting an annotation different from <u>default</u>. The overlooking could be a result of annotator fatigue, which may be attributed to length and/or complexity of the radiology report. Another data entry issue appears with derived lexical targets, which may be attributed to differences in how the lexical targets are interrelated, and hence, their derivation is differently inferred. In other cases, differences in interpretation of the radiology report were the basis for annotator disagreement. For example, “mixed density lesion” was attributed to *hypodensity* in one case and *hyperdensity* in another. There was also a difference in whether “parenchymal contusion” was considered an *intraparenchymal hemorrhage*. However, the differences between the annotators was minimal and therefore provided a valid gold standard to develop and validate tbiExtractor.

Standard assessment metrics for evaluating tbiExtractor were exceptionally high, demonstrating the utility of the algorithm for extracting accurate clinical conditions relevant to TBI research. Additionally, 93% of radiology reports in the validation dataset were accurate for over 85% of the lexical targets. While the errors from tbiExtractor on the validation dataset were minimal, there are a few cases worth exploring. First, regular expressions are unable to handle complex syntax and semantics to select the lexical target. One particularly error-prone case was *facial fracture*. Often, radiology reports with *facial fractures* are lengthy and involve compound sentence structures, which are missed by the regular expressions and span pruning. Second, there were several cases where the lexical modifier was absent or at a distance further away than another lexical modifier. For example, “cerebellar volume loss” would indicate *atrophy* is PRESENT, but with this sentence, there is no lexical modifier available and therefore would result in a <u>default</u> lexical modifier of ABSENT. Third, there were cases where derived lexical targets were not accurately annotated by tbiExtractor. After reviewing these errors, many of them were the result of ambiguous reports where “smart-phrases” had not been updated by the radiologist. These “smart-phrases” are made available in EMR systems to provide structured text statements that can easily be programmed for rapid reporting of results. For example, the sentence “there is no evidence of intracranial hemorrhage, mass effect, midline shift or abnormal extraaxial fluid collection” was frequently the first sentence in the radiology reports. This “smart-phrase” provides valuable information, however, if it is not updated, say if later in the radiology report a *subdural hemorrhage* is reported, then tbiExtractor is unable to distinguish this and annotates *extraaxial fluid collection* to be ABSENT. Further examination of these errors is an avenue for future research that may aid in optimizing tbiExtractor.

While tbiExtractor is a valuable algorithm with high performance metrics, there are limitations to its design. The dataset used for this study was from a single institution which limits the style of radiology reports and decreases heterogeneity in the sample. Furthermore, the dataset was limited in size as there were only two annotators available for annotation. In addition, there were data entry issues from extracting the radiology report from the EMRs. For example, a subsequent radiology report was used instead of the admission. Lastly, the only scan considered in this dataset is the admission non-contrast head CT. With the nature of TBIs, some visible pathologies are only seen on follow-up CTs and would be missed on initial imaging.

## CONCLUSION

tbiExtractor was developed to automate the extraction of TBI common data elements from radiology reports. Using two annotators as the gold standard, tbiExtractor displayed high sensitivity and specificity. Findings also showed high equivalence in annotations between annotators and tbiExtractor. Additionally, the time it took tbiExtractor to extract information from the radiology reports was significantly less than the time it took annotators to complete the same task. In conclusion, tbiExtractor is a highly sensitive algorithm for extracting clinical conditions of interest in TBI by providing a structured summary of their status from the radiology report. This algorithm can be used to stratify subjects into severity-groups based on visible damage by partitioning the annotations of the pertinent clinical conditions on a non-contrast head CT report.

## COMPETING INTERESTS

None

### ACKNOWLEDGEMENTS

None

